# Continuous, Low Latency Estimation of the Size and Shape of Single Proteins from Real-Time Nanopore Data

**DOI:** 10.1101/2025.07.07.663422

**Authors:** Yuanjie Li, Cuifeng Ying, Michael Mayer

## Abstract

Existing approaches for nanopore sensing typically analyze resistive pulses from the translocation of individual proteins through the nanopore after completing the experiment. This approach foregoes instantaneous protein identification and precludes real-time experimental control. Here, we introduce a method for the analysis of real-time nanopore data capable of characterizing the size and approximate shape of proteins within a millisecond response time during data acquisition. The implemented real-time Two-Sliding Window (TSW) peak detection algorithm makes it possible, for the first time, to filter data, process baselines, and extract resistive pulse information during nanopore recordings using a stream of single data points. This approach achieves a computational throughput of 40 MB/s on a 1.8 GHz laptop CPU. We compared the accuracy of dwell time determination of the TSW algorithm with an established offline threshold searching (TS) algorithm, using simulated resistive pulses. The TSW algorithm accurately extracted events with dwell times greater than 1.5 times the reciprocal of the system’s cutoff frequency, *1.5 × (fc)*^−1^. Moreover, the TSW algorithm effectively determined the dwell time of resistive pulses with large intra-event modulation, which is crucial for resolving non-spherical protein-induced fluctuations. Finally, we verify experimentally that integrating the TSW algorithm into the data acquisition process makes it possible to determine the approximate shape and volume of proteins within low-millisecond response times. This real-time data analysis approach potentially provides instant feedback control for manipulating single proteins during or immediately after translocation. This analysis also improves data storage efficiency during recordings with a high sampling rate.

## Introduction

Nanopore-based resistive pulse sensing, inspired by the principle of the Coulter Counter for blood cell counting,^1^ has shown remarkable progress in the characterization of unlabeled proteins in solution.^2–11^ Briefly, a thin insulating membrane separates two compartments of electrolyte solution, with a single nanopore serving as the only connection. Two Ag/AgCl electrodes apply a constant potential difference between the two reservoirs and measure the ionic current through the nanopore via a low-noise amplifier.^12, 13^ The passage of an insulating particle through the nanopore transiently replaces the electrolyte solution, producing a resistive pulse. These resistive pulses, also called events, can reveal various parameters of the transiting particles, including size, shape, charge, dipole moment, and rotational diffusion coefficient.^2, 14–17^

Analyzing resistive pulse data usually consists of four steps: correction for drifts in the recorded current baseline, low-pass data filtering, peak detection, and peak analysis.^18–26^ Low pass-filters, such as the Gaussian filter^3, 4^ or Butterworth filter^27–29^, are usually employed to remove the high-frequency noise. Although peak detection methods vary between research groups, they generally rely on threshold algorithms to extract events from filtered data.^18, 19, 21, 24, 30^ A common approach uses a threshold of five times the standard deviation of the recorded current baseline, also called root mean square (RMS) to distinguish resistive pulses with a 99.99994% confidence level from baseline fluctuations due to random noise.^18^ This approach starts with baseline detection using a moving average of over 100 to 30000 data points to account for long-term drift and signal fluctuations.^18–21, 24^ Several peak detection methods have been developed to improve the accuracy of event determination. Plesa et al.^19^ developed an iterative baseline approach to refine the baseline. Gu et al.^20^ introduced a second-order differential-based calibration (DBC) method to determine the dwell time and magnitude of blockades of resistive pulses. Furthermore, various fitting methods, such as Adaptive Time-Series Analysis (ADEPT),^21^ Cumulative Sum (CUSUM),^21, 31^ AutoStepfinder,^32^ and Rissanen’s Minimum Description Length (MDL)^33^ have been proposed or explored to analyze resistive pulses with short and long residence times or to detect step signals.

These established data analysis methods have in common that they typically require data to be collected over extended periods prior to analysis, thereby foregoing the possibility of real-time feedback for experimental control during recording. For example, estimating a protein’s shape requires resistive pulses with extended duration to sample various orientations. For instance, at a bandwidth of ∼50 kHz, we showed that resistive pulses should last for at least 150 µs to increase the probability that several protein orientations can be sampled.^3^ Without knowing the number of captured resistive pulses that exceed this minimum-duration threshold during a recording, it is challenging to decide when to stop the experiment.^34^ Wang et al.^22^ developed an adapted finite impulse response filter for real-time threshold searching during data acquisition. However, the threshold searching algorithm usually required additional baseline correction and resistive pulse fitting steps to accurately extract the duration and amplitude of current blockades. These processing steps are not only time-consuming but also dependent on contextual data collected during the experiment. The high sampling rates during nanopore recordings, such as 500 kHz, and the recently developed parallel recordings^35^ make low-latency, high-precision data analysis particularly challenging. Therefore, developing a robust, real-time processing method that enables accurate event detection during nanopore experiments is needed.

Here, we adopt a two-sliding window (TSW) algorithm to detect resistive pulses from translocation events accurately and instantly, making it possible to characterize the volume and approximate shape of proteins while they are still inside the nanopore or shortly thereafter. This TSW approach applies the 5×RMS and the *z*-test^30^ criteria to detect the start and end of resistive pulses and, hence, its duration and amplitude. We implemented the approach with incremental one-pass algorithms^36^, which enable real-time detection of resistive pulses with a time cost for computation linear to the number of data points. Using simulated square pulses, we demonstrate that TSW algorithm performs as well as the established threshold searching algorithm (TS) for the detection of resistive pulses and provides improved accuracy in determining dwell times of long resistive pulses or resistive pulses with large intra-event current modulations. We implemented the TSW algorithm into the real-time protein characterization method and used it for experiments to determine the shape and volume of two natively folded proteins with non-spherical shape, Immunoglobulin G (IgG) and Thyroglobulin (Tg). We demonstrate that this approach achieves accurate estimates of protein volume and shape during the recording while the protein is still in the pore or less than 1 ms after. This work, therefore, introduces a robust and low-latency method to estimate the shape and volume of individual, natively folded protein molecules while they translocate through a nanopore.

## Baseline Determination and Time Requirement of the TSW Algorithm

Figure 1 presents an overview of the analysis software (called PyDAQ, see **Figure S1**) that we developed to collect data and determine real-time estimates of the size and shape of proteins. As proteins translocate through a nanopore, the ionic current exhibits resistive pulses, each corresponding to the presence of a protein in the pore (Figure 1A). These detected resistive pulses, highlighted by the green vertical lines in Figure 1B, provide the opportunity to estimate the protein’s volume and shape in real time, based on the observation that non-spherical proteins produce orientation-dependent current modulations as opposed to spherical proteins, which produce resistive pulses that do not depend on protein orientation in the electric field (Figure 1C).^2^ Figure 1D shows that as the number of detected resistive pulses from a solution of pure protein accumulates, the estimated shape and volume converge to a stable, accurate value after an accumulated resistive pulse time of 18 ms. The results demonstrate that extending the recording time of protein translocation beyond this time will bring no further benefit regarding the accuracy of volume and shape determination. This real-time feedback on protein parameters minimizes the duration of nanopore recording needed for accurate analysis, thereby improving the efficiency of sensing.

**Figure 1.**
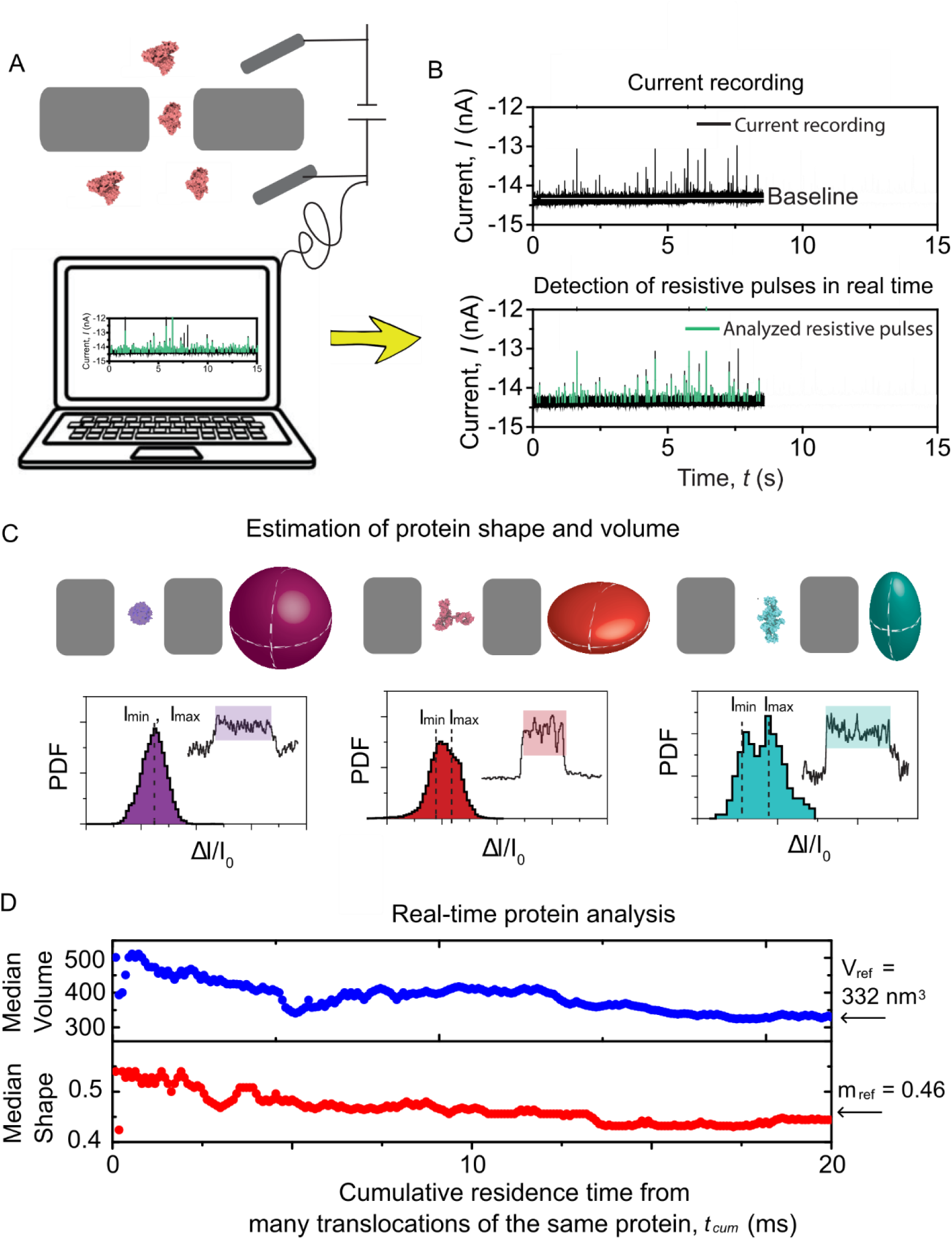
Estimating the volume and shape of proteins during nanopore data acquisition. **A.** Schematic illustration of the nanopore recording setup and of the real-time data analysis approach for protein characterization. The electrolyte contains 2 M KCl with 10 mM HEPES buffered at pH 7.4. Two Ag/AgCl electrodes apply a potential difference of –100 mV across a nanopore with a diameter of 20 nm and a length of 30 nm (with negative polarity applied to the top). **B.** Representative current recording of protein translocations through a nanopore (top), with resistive pulses detected instantaneously by the TSW algorithm as indicated by green pulses (bottom). **C**. Principle of determining shape and volume of proteins from *I_min_* and *I_max_*. Top: translocation of different proteins with their shapes represented as a sphere (streptavidin), oblate (IgG), and prolate (Tg). Bottom: Representative current pulses generated from protein translocations and their corresponding histogram, where *I_min_* and *I_max_* are the minimum and maximum current blockades of the single resistive pulse. The ratio between the magnitude of *I_min_* and *I_max_* determines the shape *m* of proteins. Here, *m* is defined as the axis ratio *b/a* of an ellipsoid with semi-axes *a, a, b*; *m < 1* corresponds to an oblate shape, while *m > 1* indicates a prolate shape. **D.** Estimation of shape (*m*) and volume (*V*) from the cumulative residence time of detected resistive pulses during recording for a protein modeled as an oblate shape IgG. The arrows represent that the estimated volume and shape stabilize at *V_ref_* = 332 nm^3^ and *m_ref_* = 0.46 after a cumulative residence time of 18 ms. Here, only the resistive pulses with dwell times greater than 150 µs were analyzed. Data were acquired at a sampling rate of 500 kHz and a bandwidth of 50 kHz.

The approach for real-time protein characterization introduced here consists of four independent processes, which are illustrated in the flowchart in Figure 2. These processes include data acquisition, low-pass filtering with event detection, data visualization with data storage, and estimation of protein size and shape. The algorithm continuously acquires the data at the maximum sampling rate of the recordings of 500 kHz and temporarily stores them in a hardware buffer with a size of 10 k samples. Simultaneously, the TSW algorithm performs low-pass filtering with a real-time Butterworth filter and peak detection on the data stream (right inset of Figure 2). Another computing process stores and visualizes the data, while a new process estimates protein size and shape according to the peak detection results of the TSW in real time.

**Figure 2.**
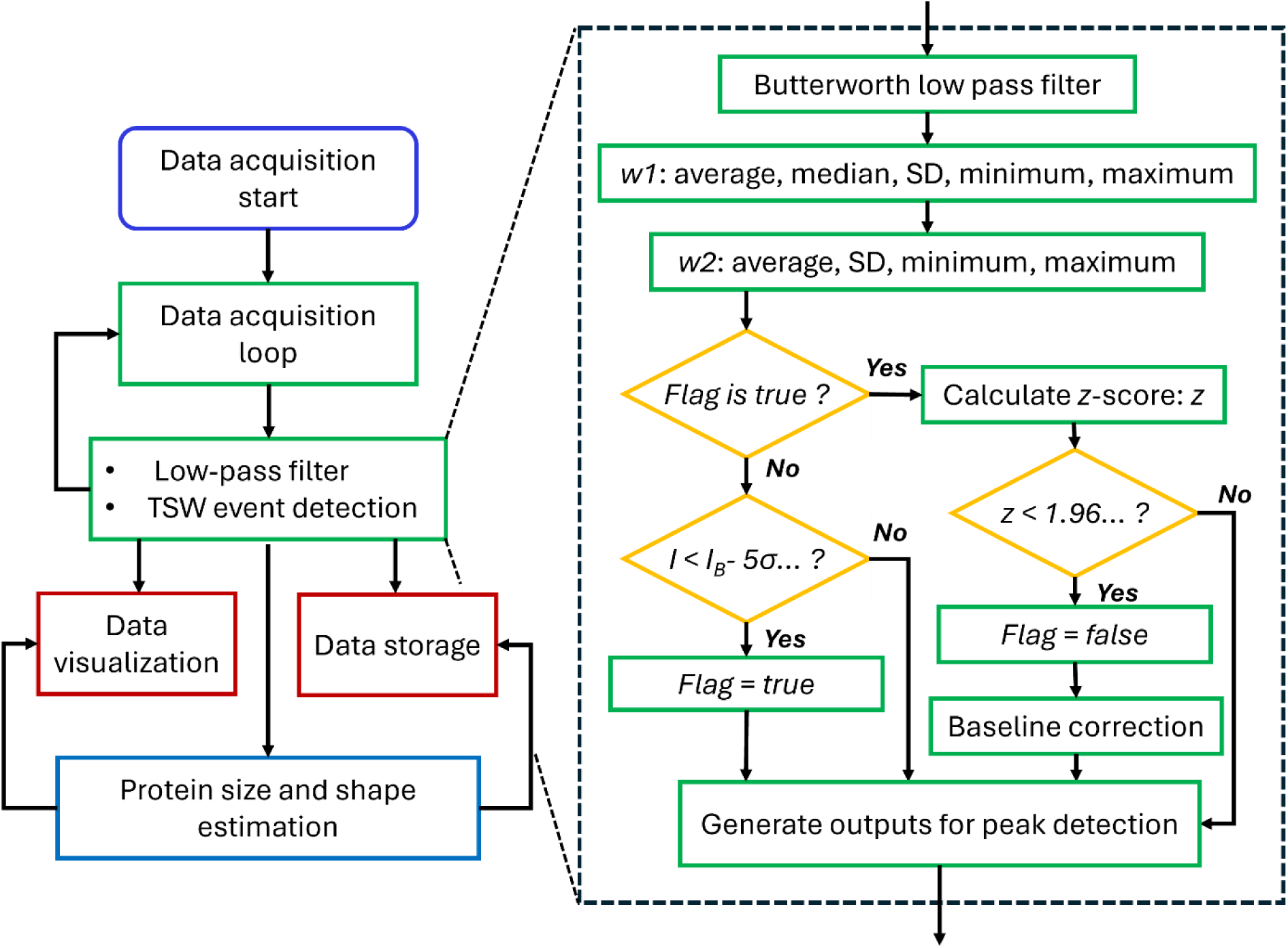
Flowchart illustrating the process of real-time determination of the size and shape of proteins from nanopore recordings. Data is collected from a data acquisition card every 20 ms in each loop. Concurrently, data are processed in real time via a low-pass Butterworth filter and the TSW algorithm (inset). Detected events are visualized in real time and stored on disk by a different task process. Concurrently, within 20 ms, an independent thread estimates the protein parameters by continuously receiving the detected resistive pulses, calculating the shape and volume, and feeding this information back to a thread for visualization and file storage.

Briefly, the TSW algorithm uses two moving windows, *w_1_,* and *w_2_*, to accurately localize the start and end of a resistive pulse, where *w_1_* is a variable-length window for estimating baseline parameters, and *w_2_* is a fixed-length window for estimating parameters of resistive pulses. These windows update the median, mean, standard deviation, minimum, and maximum value for the measured current using established incremental one-pass algorithms^36–38^ (see **Supplementary Note 1**). Figure 3A visualizes how a parameter called sigma level (*n-sigma,* red) and a parameter called *z-score* (*z*, blue) change over time during nanopore recordings. These two parameters facilitate the identification of the start and end of resistive pulses. The sigma level is the ratio between the amplitude of the resistive pulses and the standard deviation of the baseline (*n-sigma* = *(I – I_0_) / σ*). The TSW algorithm detects the start of a resistive pulse when *n-sigma* is larger than 5, i.e., *I – I_0_ >* 5*σ*, where *σ* is the moving standard deviation, *I_0_* is the average or median current, *I* is the current of pulses in window *w_1_*. Unlike the global noise used by our group and many others previously, the moving standard deviation used here can improve the adaptability of the peak detection algorithm in order to analyze unstable current baselines during experiments. To reduce the errors for the determination of the end of resistive pulses caused by single-point-based criteria, the TSW algorithm computes the *z-score* (*z*) for each data point and applies a *z*-test threshold (*z < 1.96*) to determine the end of resistive pulses, as described in **Eq 1**.

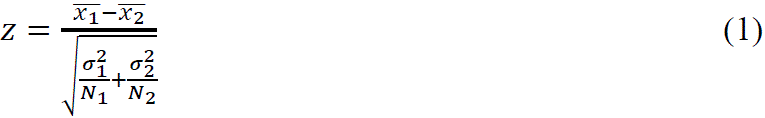

**Figure 3.**
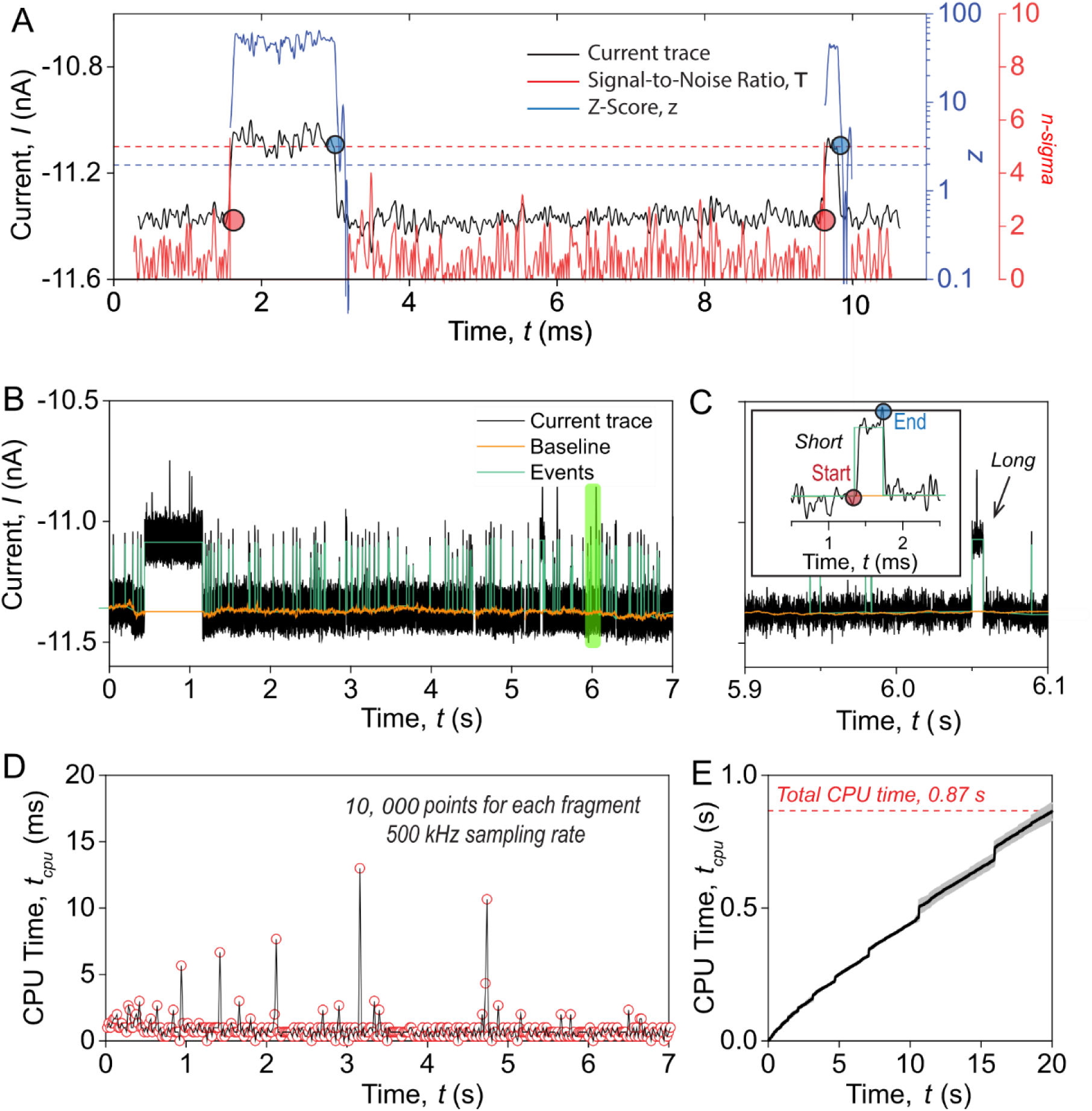
**Principle and time complexity of the TSW algorithm for real-time analysis of resistive pulses data**. **A**. Signal-to-noise ratio (red) and *z-score* (blue) as a function of time during the analysis of a resistive pulse. The red dashed line shows the reference value of *n-sigma* of 5, which has to be exceeded to detect the start of a resistive pulse. The blue dashed line shows the threshold z*-score* of 1.96 for detecting the end of a resistive pulse at a confidence level of 99.9%. **B**. Experimental current recording of resistive pulses from protein translocations through a nanopore. The orange curve represents the baseline determined using the TSW algorithm, and the green curves indicate the identified resistive pulses using the TSW algorithm. **C.** Zoomed-in region from **B** (light green shading) showing a detected short resistive pulse with start (red) and end (blue) points, along with an arrow indicating a detected long resistive pulse. **D**. Time of CPU processing during analysis as a function of data acquisition time. Red circles represent the time consumption for data updated every 20 ms, with a sampling rate of 500 kHz. Spikes in CPU time are due to the process scheduling by the operating system. **E**. Accumulated CPU time for processing a current trace with a duration of 20 seconds.

Here, *z* is the *z*-score, 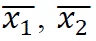 are the mean currents and *σ_1_*, *σ_2_* are the standard deviations calculated in *w_1_, w_2_*. The values of *N_1_* and *N_2_* represent the initial size in terms of the number of data points within *w_1_* and *w_2_*. Only data points with *z-score* of less than 1.96 (at the confidence level of 99.9 %) indicate the end of a resistive pulse. We also constrain the maximum and minimum values of the baseline within the window *w_2_* to remain within the range *I_0_ ± 5σ_1_* (**Figure S2**), which is useful to detect resistive pulses with non-Gaussian distribution of the blockade current, such as the current blockade from the translocation of non-spherical proteins.

The determined baseline, represented by the orange curve in Figure 3B, agrees well with the expected baseline throughout the entire recording, even in regions with high event frequency or events that last unusually long times (Figure 3C). In contrast, the baseline determined by the established threshold searching (TS) algorithm is strongly influenced by the event frequency and duration (see **Figure S3**). As opposed to the established TS with a fixed size of the moving analysis window,^3, 18, 19^ the TSW algorithm varies the window size during analysis: the window size increases or remains unchanged when there are no resistive pulses and stops updates of its size when a resistive pulse occurs. Therefore, the TSW algorithm can effectively eliminate the influence of resistive pulses on the baseline. In addition to an accurate determination of baseline, the TSW algorithm minimizes the effects of the filter rise time on the determination of dwell times by defining the start of the rising edge as the start of a resistive pulse and the start of the falling edge as the end of a resistive pulse (Figure 3C). The start and end of resistive pulses detected by the TSW algorithm are comparable to those determined by the TS algorithm (see **Figure S3**). The TSW algorithm offers two optional approaches to calculate the baseline, moving median, and moving average (see **Supplementary Note 1, algorithm1**). These two approaches for baseline determination proceed with different speeds since the analysis time of the moving average is proportional to the number of data points, *n*, but the analysis time of the moving median is proportional to *nlog (w_1_)*, where *n* is the number of points and *w_1_* is the size of the *w_1_* window. Although the moving median algorithm is slightly slower than the moving average algorithm, it provides a more accurate baseline determination due to its robustness against outliers.

The computational efficiency of peak-finding algorithms is essential for achieving instant analysis. Figure 3D plots the CPU time required to process each 20 ms iteration as the data are continuously streamed into the analysis program. For protein detection, we collect data at a high sampling rate of 500 kHz using a data acquisition (DAQ) card with a specific buffer size of 10,000 data points. When the buffer accumulates 10,000 data points (every 20 ms), the TSW algorithm, instantly, processes the collected data. It is important to note that although we analyzed the data stream in each 20 ms loop, the TSW algorithm analyzes the data streams by receiving a single point as input, and therefore, the analysis proceeds as each data point is acquired and, hence, is made available and processed in real-time. Moreover, smaller buffer sizes than 10,000 data points are feasible when using a real-time operating system such as FreeROTS^39^. The results in Figure 3D demonstrate that the TSW algorithm processes every 20-ms segment of data within 1 ms on average, hence keeping up with the analysis of data while it is being recorded. Figure 3E suggests that the overall time consumption of the TSW algorithm scales linearly with the size of input data, corresponding to a time complexity of *O(n)*. For processing a current trace (i.e., 20 seconds) on a laptop CPU with a base clock frequency of 1.8 GHz, the TSW algorithm only takes 0.87 s, corresponding to an analysis speed of 40 MB/s for the data stream. We note that changes in baseline window size, *w_1_*, do not affect the processing speed of the TSW algorithm (see **Figure S6)**. The results reveal that the time consumption of the TSW algorithm is influenced solely by the number of recording data with no significant impact from other parameter settings.

## Comparison of the TSW algorithm and the threshold searching algorithm for the detection of resistive pulses

To evaluate the performance of the TSW algorithm for the detection of resistive pulses (see Figure 4) and to compare it with the commonly used TS algorithm,^18^ we simulated various current blockades by adding Gaussian noise with ideal square wave pulses. We generated these blockades at a frequency of 1 Hz and a sampling rate of 500 kHz, followed by filtering with a Gaussian low-pass filter with a cutoff frequency of 50 kHz. We compare the performance of resistive pulse detection for two different scenarios: various dwell times with the same baseline noise and constant standard deviation of intra-event modulation (Figure 4A), versus constant dwell time and baseline noise but various standard deviation of intra-event modulations (Figure 4B).

**Figure 4.**
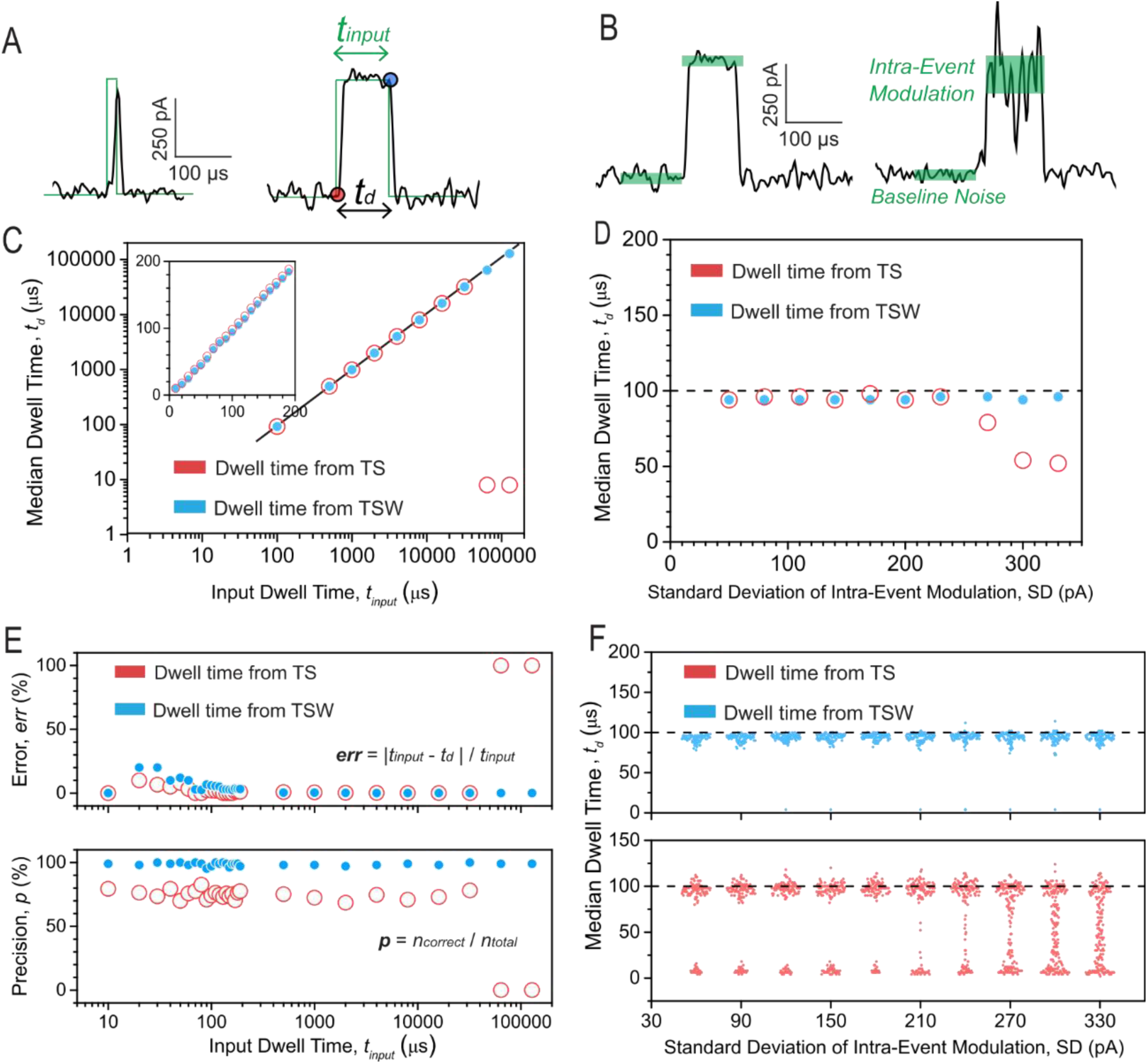
Comparison of the TSW and TS algorithms for dwell time analysis of resistive pulses. **A**. Ideal square waves of varying pulse width overlapped with their generated blockades (black). The green line represents the ideal square wave with input dwell times ranging from 10 μs to 128 ms and a current amplitude of 500 pA. The black curve represents the generated blockades by adding Gaussian noise with a standard deviation of 50 pA to the ideal square wave. The dwell time *t_d_* is defined between the detected start and end points (red and blue), identified by the TSW or TS algorithms. **B**. Generated blockades with different intra-event modulations. The green shaded regions indicate the current baseline with a noise of 50 pA and the standard deviation of intra-event modulations varying from 50 pA to 330 pA. All the generated blockades have a sampling rate of 500 kHz, followed by low-pass filtering with a Gaussian filter with a cutoff frequency of 50 kHz. **C**. Median dwell time evaluated by the TSW (blue) and TS (red) algorithms as a function of input dwell time. The black line represents perfect agreement of the dwell time. Inset: The input dwell time of generated blockades varies from 10 μs to 200 μs. **D.** Median dwell time evaluated by the TSW (blue) and TS (red) algorithms as a function of standard deviation of intra-event modulations. The black dashed line represents the input dwell time of 100 μs for all the generated blockades. **E.** Relative error and precision of detected blockades from the TSW (blue) and TS (red). The parameter *n_total_* represents the overall number of events that are detected. The parameter *n_correct_* represents the detected events that belong to the ideal square waves. **F.** Box chart of dwell time evaluated by the TSW (blue) and TS (red) from the generated blockades of varying standard deviation of intra-event modulations. The black dashed line represents the input dwell time of 100 μs for all the generated blockades.

When the standard deviation of intra-event modulation is the same as that of baseline noise, both the TSW and TS algorithms evaluate the dwell time accurately, as shown **in** Figure 4C. However, the TS algorithm fails to detect long blockades with dwell times greater than 60 ms, whereas the TSW algorithm consistently detects these long events with accurate dwell times. Figure 4E shows that compared to the TS algorithm, the TSW algorithm exhibits a 10% higher relative error of dwell time for the resistive pulses with dwell times shorter than 30 μs, indicating that the TSW algorithm may be slightly less reliable than the TS algorithm for resistive pulses with *t_d_* < 1.5 × (*fc)*^−1^. This error arises from noise fluctuations, which are challenging to distinguish from short events. The TS algorithm employs an additional fitting step on the rising edge and falling edge to accurately recover the entire short resistive pulses. However, as long as *t_d_* > 1.5 × (*fc)*^−1^, the errors of the TSW algorithm are comparable to those of the TS algorithm. Additionally, the precision of the TSW algorithm is, on average, 25% better than that of the TS algorithm (Figure 4E**, bottom**). The results demonstrate the robustness of the TSW algorithm in handling noise, which we attribute to the benefit conferred by the *z-test* criterion on determining the end of resistive pulses.

Protein translocation events exhibit significant fluctuations within each resistive pulse, so called intra-event modulations, as shown in Figure 1C. These modulations arise from the rotation of particles with non-spherical shape in a nanopore and can be used to estimate the shape of proteins.^2^ To verify the effectiveness of the TSW algorithm in handling resistive pulses with large intra-event modulation, we generated a series of blockades with the standard deviation of intra-event modulations ranging from 50 pA to 350 pA (Figure 4B). Figure 4D shows that as intra-event modulations increase, the median dwell time determined by the TSW algorithm closely aligns with the input dwell time of 100 μs. In contrast, the dwell time determined by the TS algorithm gradually deviates by approximately 50% from the reference value when the standard deviation of intra-event modulation exceeds 250 pA. The box plot in Figure 4F further illustrates that the dwell times detected by the TSW algorithm remain consistent with the reference dwell time of 100 μs across different standard deviations of intra-event modulations. In contrast, the detected dwell time evaluated by the TS algorithm exhibits a broader distribution than the dwell time distribution evaluated by the TSW algorithm, and this distribution becomes further distorted with increasing standard deviation of intra-event modulation. The TS algorithm struggles to identify the correct duration of resistive pulses in the presence of large intra-event modulations, as these modulations significantly diminish the determination of event endpoints the single-point criterion of the TS algorithm. In contrast, the TSW algorithm effectively extracts the complete blockades with these intra-event modulations by using a *z-test*. This capability is crucial for protein characterization, as it enables accurate extraction of all current modulations induced by protein rotations or conformational changes.^40^

Next, we explored the performance of the TS and TSW algorithms on simulated protein translocation data (**Supplementary Note 2, Figure S5, Figure S7)**. The TSW algorithm shows a lower standard deviation of 0.04% on determining *ΔI/I_0_* values compared to 0.08% when using the TS algorithm. We evaluated the agreement between reference dwell times and determined dwell times as detected by TS and TSW using the sum of squared residuals (SSR). The SSR of the TS and TSW algorithms were 4.8 × 10^−6^ and 17.0 × 10^−6^, respectively, suggesting that the TSW algorithm has a slightly larger error in dwell time determination for protein translocation data. This increased error is due to the reduced accuracy of the TSW algorithm when determining dwell times of very short resistive pulses with dwell times close to the bandwidth limitation (*t_d_ ≤ 1.5* × (*fc)*^−1^). However, this limitation does not affect the characterization of individual proteins, as only events with dwell times greater than *2.5* × (*fc)*^−1^ were considered in the estimation of protein shape and volume.^2, 3^ The results reveal that the detection of resistive pulses by the TSW algorithm performs as well as the previously employed TS algorithm for the events with dwell time greater than *1.5* × (*fc)*^−1^, while providing a 0.04% improvement in the standard deviation of *ΔI/I_0_* values compared to the TS algorithm under these conditions.

## Instantaneous determination of the size and shape of single proteins

With instantaneous and accurate identification of resistive pulses from the incoming stream of data, we explored the application of the TSW algorithm for determining the shape and volume of proteins from experiments. Previously, we demonstrated the estimation of shape, volume, and other parameters of proteins from their free translocation through nanopores.^3^ Due to bandwidth limitations when applying a low-pass filter with a cutoff frequency of 50 kHz, only resistive pulses longer than 50 µs can be used for population-based analysis^2^, and only resistive pulses longer than 150 µs are suitable for the analysis of protein shape from individual resistive pulses. Obtaining a sufficient number of long events for precise protein characterization using conventional offline peak detection methods remains challenging, and the required recording time is typically estimated empirically to be on the order of several minutes. The TSW algorithm offers the flexibility to terminate data acquisition once a sufficient number of analyzable events has been collected.

To test the usefulness of the TSW algorithm for real-time estimation of protein shape and volume, we conducted translocation experiments with Immunoglobulin G (IgG, oblate shape, *m* = 0.4) and thyroglobulin (Tg, prolate shape, *m* = 2.1) using solid-state nanopores with diameters of 25 and 35 nm and with a coating of PMOXA polymer. Following the flowchart in Figure 2, we detected translocation events and determined the shapes and volumes of the two separate proteins in real time using PyDaq (Figure 5). Figure 5A **and 5B** depict the current recording of IgG and Tg translocating through nanopores, respectively, with detected resistive pulses indicated by orange dots. The passage of individual IgG or Tg proteins resulted in characteristic resistive pulses, as indicated at bottom of Figure 5A and **5B**. To estimate the shape and volume of the proteins, we fit the cumulative distribution of the current within individual resistive pulses whose *t_d_* values exceeded 150 μs to a convolution model developed by Yusko et al.^2^ Figures 5C **and 5D** display the estimated length-to-diameter ratio (*m*) and volume (*V*) for IgG and Tg as a function of the cumulative residence time of detected resistive pulses. When the cumulative residence time of resistive pulses is less than 20 ms, the determined shape and volume of IgG fluctuate, with *m* between 0.2 and 0.8, and *V* between 200 and 360 nm^3^. These values then converge to *m* = 0.5 and *V* = 350 nm^3^ (*m_ref_* = 0.46, *V_ref_* = 332 nm^3^) once the cumulative number of resistive pulses exceeds 20 ms (Figure 5C). The estimated shapes and volumes of Tg show larger uncertainty, with *m* ranging from 1.2 to 3.1 and *V* ranging from 500 to 3000 nm³ during the initial 40 ms. Subsequently, these estimates converge to a value of *m* = 2.1 and *V* = 1500 nm^3^, which is close to the expected reference values of *m_ref_* = 1.9 and *V_ref_* = 1247 nm^3^. (Figure 5D). These results highlight that the estimated shape and volume remain unchanged over time once the cumulative residence time reaches a specific threshold. The TSW algorithm demonstrates its ability to extract events during data acquisition and is compatible with protein characterization methods for shape and volume estimation during experiments.

**Figure 5.**
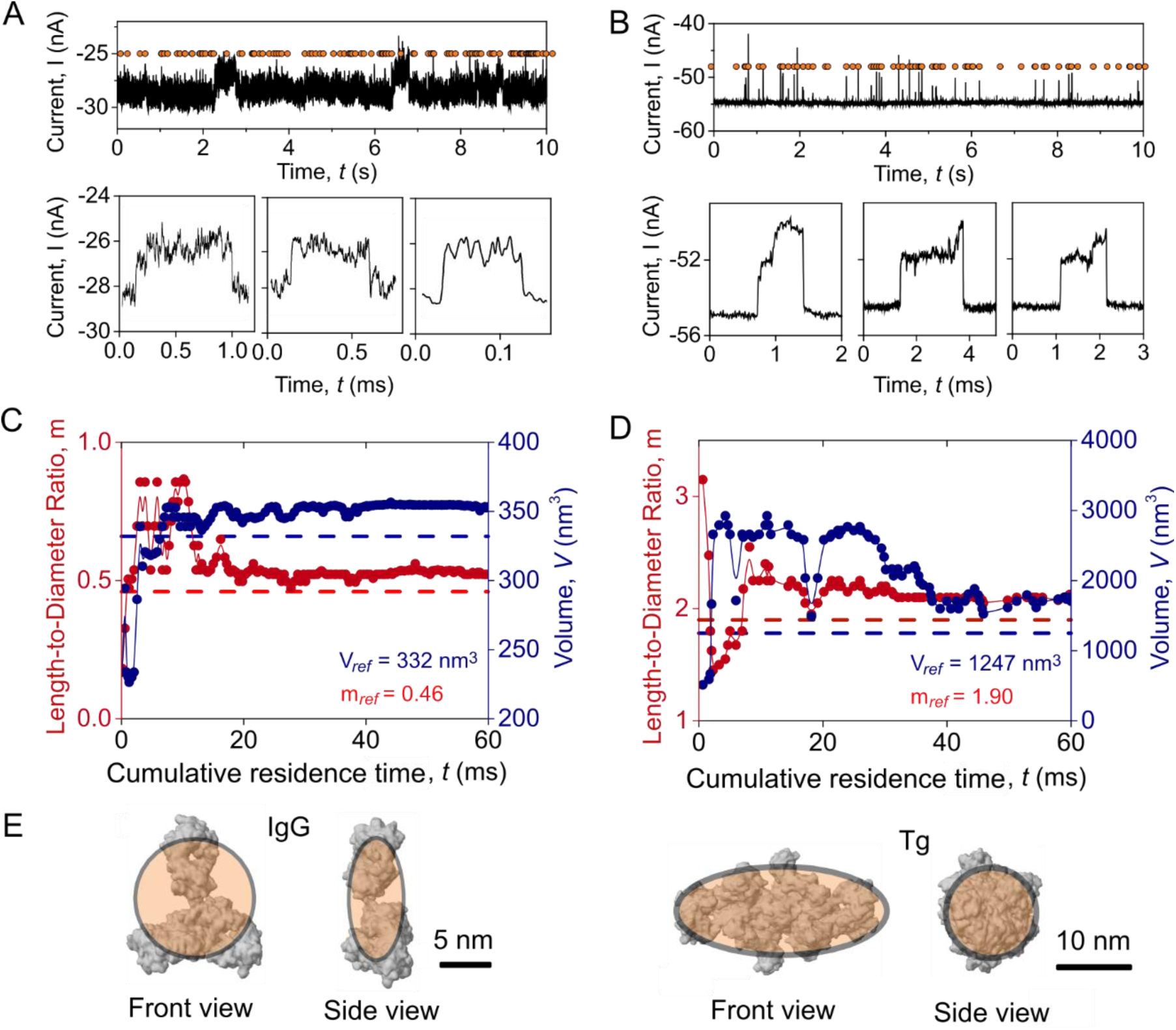
Real-time estimation of protein length-to-diameter ratio and volume from nanopore recordings using the TSW algorithm. A, **B.** Representative current recordings of IgG (**A**) or Tg (**B**) translocations through nanopores, along with three individual resistive pulse examples. Orange circles represent resistive pulses detected by the TSW algorithm. **C, D**. Real-time measurements of the length-to-diameter ratio *m*, and volume *V* for IgG (**C**) and Tg (**D**), plotted as a function of cumulative residence time. The blue and red dotted lines mark the reference length-to-diameter ratio *m* (IgG = 0.46, Tg = 1.9) and volume *V* (IgG = 332 nm^3^, Tg = 1247 nm^3^) as estimated by minimum volume enclosed ellipsoids fitting based on solvent-excluded surface. Only the resistive pulses with dwell times of greater than 150 μs were included in the cumulative residence time and used for the shape and volume estimation. The overall recording times were 80 s for IgG and 210 s for Tg. **E.** Crystal structures of IgG (PDB: 1HZH) and Tg (PDB: 6SCJ) overlaid with their best-fit ellipsoidal models (orange shadow).

## Conclusion

In conclusion, the work presented here introduces a real-time data analysis method for characterizing the shape and volume of proteins during data acquisition, providing instantaneous results within a few milliseconds and therefore before the next resistive pulse is recorded. The two sliding window (TSW) algorithm introduced here for analysis of resistive pulses enables instant event analysis with a constant speed of 40 MB/s using a CPU of 1.8 GHz base clock frequency, while the data acquisition had a speed of 2 MB/s at 500 kHz sampling rate. Using a variable-length window, the TSW algorithm determines the baseline in real time and remains unaffected by high event frequency, large intra-event modulations, or long-lasting resistive pulses. This algorithm provides more accurate baseline values compared to those determined by the threshold searching (TS) algorithm. Additionally, we incorporate the *z*-test criterion to determine the end of resistive pulses in the TSW algorithm, enhancing the robustness of peak detection for resistive pulses with large current modulations. Using the TSW algorithm, we developed an instantaneous protein characterization method enabling real-time estimation of the shape and volume of proteins during nanopore experiments. This method, hence, has the potential for real-time feedback control and enhanced storage efficiency, and may, in the future, enable sorting of individual molecules based on instantaneous analysis of the incoming data stream during their translocation through a nanopore.

## Materials and Methods

### Materials

The polymer poly(acrylamide)-g-PEG-PMOXA for coating solid-state nanopores was obtained from SuSoS AG, Switzerland. Thyroglobulin, human (T6830-1MG) was purchased from Sigma-Aldrich, USA. Immunoglobulin G, human plasma (340-21) was purchased from Lee Biosolutions, USA. Anotop 10 mm Syringe Filters with 20 nm pore size (6809-1002) were purchased from Fisher Scientific. Nanopores with a diameter of 25 nm or 35 nm in the freestanding 30-nm-thick silicon nitride membrane were ordered from Norcada Inc., Canada.

### Surface Coating

Nanopores were coated using Poly(acrylamide)-g-PEG-PMOXA, as described by Awasthi et al.^34^ After oxygen plasma cleaning (Diener Nano, 30 % power, O2 atmosphere, 0.3 mbar) for 30 s on each side, the nanopore chips were immersed in 1 mg/mL Poly(acrylamide)-g-PEG-PMOXA polymer in a buffer solution (1 mM HEPES pH 7.4) and incubated for 1 h at room temperature. These chips were rinsed with ultrapure water three times and dried under a stream of *N_2_* gas.

### Electrical Recordings

We used Ag/AgCl pellet electrodes (Warner Instruments) to monitor the ionic currents during nanopore experiments using patch-clamp amplifier (AxonPatch 200B, Molecular Devices) under voltage clamp mode with a 100 kHz lowpass Bessel filter. The data were acquired using a data acquisition card (NI PCI 6281, National Instrument) and our custom control software, PyDAQ, at a sampling rate of 500 kHz. Nanopore resistance was calculated by the slope of the ionic current at various applied voltages in the range of ±0.5 V. All experiments were performed in a recording buffer consisting of 2.0 M KCl, 10 mM HEPES, pH 7.4. This recording buffer was filtered by a membrane filter with a pore size of 20 nm prior to use.

### Determination of Protein Volume and Shape

We converted the PBD structure of proteins to the solvent-excluded surface^41^ using our custom depth-first search algorithm. Protein volumes were then calculated based on the solvents-excluded surfaces, employing a water probe with a diameter of 0.28 nm. To estimate the ellipsoid parameters (*a, a, b*), we use the minimum volume enclosing ellipsoids (MVEE) algorithm^42^, which optimizes the semi-axes based on the solvent-excluded surface to determine the ellipsoid shape. We provided an executable file to perform the shape and volume fitting for the PDB file upon request.

### Computing Platform and Software

The TS algorithm is provided from a MATLAB script based on the method proposed by Pedone et al.^18^ We wrote the TSW algorithm with *C++* and compiled it to a dynamic link library for usage in Python-based data acquisition software (PyDAQ, **Figure S1**). The control software, PyDAQ, was written with Python based on a Python library *nidaqmx* (National Instruments) and has been highly optimized for visualization. It includes a fast real-time algorithm for down-sampling that is integrated into the TSW algorithm, as well as image reuses, and multithread rendering. We implemented the simulation of ellipsoid translocations with *C++* as discussed in **Supplementary Note 2**. The simulation program and the TSW algorithm were tested on a laptop equipped with a 1.8 GHz Intel Core i5 CPU and 16 GB of memory. The associated software is available on our GitHub or upon request.

## Acknowledgments

M.M. acknowledges financial support from the Swiss National Science Foundation (Grant number: 200020_197239) and from the Adolphe Merkle Foundation. Y.J.L. thank Andela Vracar for constructive feedback on the writing of the manuscript.

