## Supplementary Information for "Continuous, Low Latency Estimation of the Size and Shape of Single Proteins from Real-Time Nanopore Data"

### Supplementary Note 1. Algorithms for calculating mean, median, standard deviation, minimum, and maximum in the sliding windows

*Sliding mean and standard deviation*

The TSW algorithm calculates the sliding mean using the following equation at each iteration of the program.

$$\overline{x_n} = \overline{x_{n-1}} + \frac{x_n - x_{n-k}}{k} \qquad (1)$$

Here, the $\overline{x_n}$ is the average current in the sliding window at the position of *n*, and $x_n$ is the current at the position of *n*. The *n* is the *nth* point received from the input. The parameter *k* is the size of the sliding window.

The TSW algorithm calculates the sliding standard deviation using the following equation at each iteration of the program.

$$d_n^2 = d_{n-1}^2 + (x_n - x_{n-k})(x_n + x_{n-k} - \overline{x_n} - \overline{x_{n-1}}) \qquad (2)$$



This manuscript has been submitted to Analytical Chemistry.

Here, the $d_n^2$ is the non-normalized variance of the current in the sliding window at the position of *n*.

*Sliding median*

The TSW algorithm uses the balanced binary tree (*i.e., Red-Black Tree*)[1] as the sliding window to calculate the sliding median value on the current trace, as shown in **algorithm 1**.

Algorithm 1. Sliding Median

```
Input: timeSeries; Output: medianValues
Initialize redBlackTree as an empty set; Initialize medianValues as an empty array.
For i from 0 to length(timeSeries) – 1
        Insert timeSeries[i] into redBlackTree
        If the size of redBlackTree is greater than windowSize
                Delete redBlackTree[i - windowsize]
        Else
                medianValues[i] = redBlackTree[windowSize/2]
Return
```

*Sliding minimum and maximum*

The TSW algorithm uses *dequeue* as the sliding window to calculate the sliding *minimum*, and *maximum* value on the current trace, as shown in **algorithm 2**.

Algorithm 2. Sliding Minimum, Maximum

```
Input: timeSeries; Output: minValues, maxValues
Initialize maxque and minque as empty double-ended queues
Initialize medianValues as an empty array
For i from 0 to length(timeSeries) – 1
        While not maxque.empty() and maxque.front()<= i - windowSize
                maxque.pop_front()
        While not minque.empty() and minque.front()<= i - windowSize
                minque.pop_front()
        While not maxque.empty() and timeSeries[maxque.back()]<=timeSeries[i]
                maxque.pop_back()
        While not minque.empty() and timeSeries[minque.back()]>=timeSeries[i]
                minque.pop_back()
        maxque.push_back(i); minque.push_back(i)
        maxValues.append(timeSeries[maxque.front()])
        minValues.append(timeSeries[minque.front()])
Return
```

### Supplementary Note 2. Simulation of protein translocation through nanopores

*Simulation of intra-event current modulations*

The fundamental principle of nanopore-based characterization of protein volume and shape entails deciphering the rotation of a single non-spherical particle as it translocates through a cylindrical nanopore with uniform electric field. The induced current modulation resulting from this rotation can be used to determine the protein shape and volume. The concept draws upon the work of Golibersuch[2], who demonstrated both the theoretical concept and experimental techniques for determining the geometry of red blood cells, which exhibit an oblate shape.[3] As red blood cells pass through and rotate within an electrolyte-filled microchannel, they distort the electric field, resulting in changes in the ionic current that are directly related to the shape of blood cells. Assuming the object undergoes random rotation along one axis, the corresponding electrical shape factor, $\gamma$, can be expressed by Equation 3. Here, $\gamma_v$ and $\gamma_h$ are shape factors when the protein's singleton axis (the normal vector of the circle plane) is oriented perpendicular to or in parallel with the nanopore channel, respectively, and $\theta$ is the angle between the protein's singleton axis and nanopore cross-section plane.

$$\gamma = \gamma_v + (\gamma_h - \gamma_v)cos^2(\theta) \qquad (3)$$

Proteins undergo frequent collisions by solvent molecules in solutions, resulting in Brownian motion.[4, 5] Brownian motion was elucidated by Einstein with Equation 4. Here, $D$ $(m^2/s)$ is the diffusion coefficient, $\eta$ (Pa·s) is the viscosity of the solution, $r$ $(m)$ is the radius of the particles, $k_B$ (J/K) is the Boltzmann constant and $T$ $(K)$ is the absolute temperature.

$$D = \frac{k_B T}{6\pi\eta r} \qquad (4)$$



This manuscript has been submitted to Analytical Chemistry.

The diffusion coefficient, *D*, is related to the hydrodynamic radius of the particle, $r_h$.[3] Assuming that the protein is an ellipsoid, the rotation of a protein can be described by the rotational diffusion coefficient:

$$D_r = \frac{3kT}{16\pi\eta a^3} / (\frac{1 - \frac{1}{m^4}}{\left(2 - \frac{1}{m^2}\right) G(m) - 1}) \qquad (5)$$

Here, the shape *m (unitless)* is the ratio of the two axes of the ellipsoid, and *a (m)* is the radius of the short axis. With this rotational diffusion coefficient, we can calculate the angular change of the protein in one dimension as a result of rotational diffusion.

$$\Delta\theta = \sqrt{2D_r\Delta t} \qquad (6)$$

Here, $\theta$ represents the angle between the vector of the singleton axis and the plane of the nanopore. Protein orientation in the nanopore affects the electric field distribution, as previously described in Equation 1 for the case of 2D free rotation.[4] In the case of 3D rotation of an ellipsoid of ratio with axes (*a*, *a*, *b*) with two independent axes, the shape factor can be described as[3]:

$$\gamma = \gamma_v + (\gamma_h - \gamma_v)\,(cos(\theta_x)\cos(\theta_y))^2 \qquad (7)$$

We simulate the current induced by protein orientation using an angular random walk approach. The probabilities of moving to $\theta_x$ and $\theta_y$ direction are equal (50 % for each) when ignoring the influence of the electric field on the dipole moment of proteins. For proteins with non-uniform surface charge, the orientation of the protein will be biased by the electric field. Equation 8 describes the probability of the direction of protein rotation. Here, $P_\pm$ is the probability of the direction of movement at the current position, *E (V/m)* is the electric field intensity, $k_B$ is the Boltzmann constant, *T* is absolute temperature, and $\theta$ represents the angle between the vector of the singleton axis and the plane of the nanopore.[4]

$$P_\pm = \frac{1}{1 + e^{\pm E\mu[\cos(\theta-\Delta\theta)-\cos(\theta+\Delta\theta)]/(2k_BT)}} \qquad (8)$$

*Simulation of the duration of resistive pulses*

The duration of each resistive pulse is simulated as two types: 1. Free translocation events, and 2. adsorption events. The simulation of translocation times starts from a biased random walk[6] of proteins from the *cis* side of the nanopore to the *trans* side. Equation 9 represents a differential form to calculate the movement of the proteins in the nanopore:

$$\Delta x = v_d \Delta t + \sqrt{2D\Delta t} * B \quad (9)$$

Here, the $\Delta x$ (m) is the updated distance for the protein during a time step $\Delta t$. $D$ is the one-dimensional diffusion coefficient of the protein, and $B$ is a random number from a standard Normal distribution. The drift velocity $v_d$ is given by $v_d = u_e E$, where $E$ is the electric field and $u_e$ is the electrophoretic mobility. According to Smoluchowski theory[7], the electrophoretic mobility is defined as $u_e = \varepsilon_r \varepsilon_0 \zeta / \eta$ ($m^2 \cdot s/V$). Adsorption of proteins to the pore wall can occur during the process of translocation. We model the adsorption process using a Markovian reaction, which reveals the distribution of dwell times of adsorption or dissociation, $t_{on}$, or $t_{off}$ by sampling from an exponential distribution.[8]

$$p(t|k_i) = k_i e^{-k_i t} \quad (10)$$

Here, $p(t \mid k_i)$ is the probability density function of lasting time, $t$, in the *ith* state, given the stochastic rate constant, $k_i$, for dissociation. **Figure S4** represents the model of protein adsorption during translocation. We calculate the dwell time of free translocation, $t_d$, from the iteration steps using Equation 9 until the proteins leave the channel of the pore. Then, we generate a random number, $t_{on}$, from the PDF distribution in Equation 8 given $k_{on}$ to determine whether the adsorption occurs by $t_{on} < t$. If the adsorption occurs, the $t_{off}$ generated from the PDF in Equation 10 with giving $k_{off}$ will be the new duration time induced by adsorption.

The event frequency is simply described by an exponential distribution, which has a parameter capture rate constant of $k_f$ ($s^{-1}$). Besides, we convolve a white noise on the simulation data to simulate the recording noise in nanopore experiments. **Figure S5** represents an example of



This manuscript has been submitted to Analytical Chemistry.

simulated current as a function of time, showing signals similar to experimental data, including resistive pulses with both short and long dwell times.

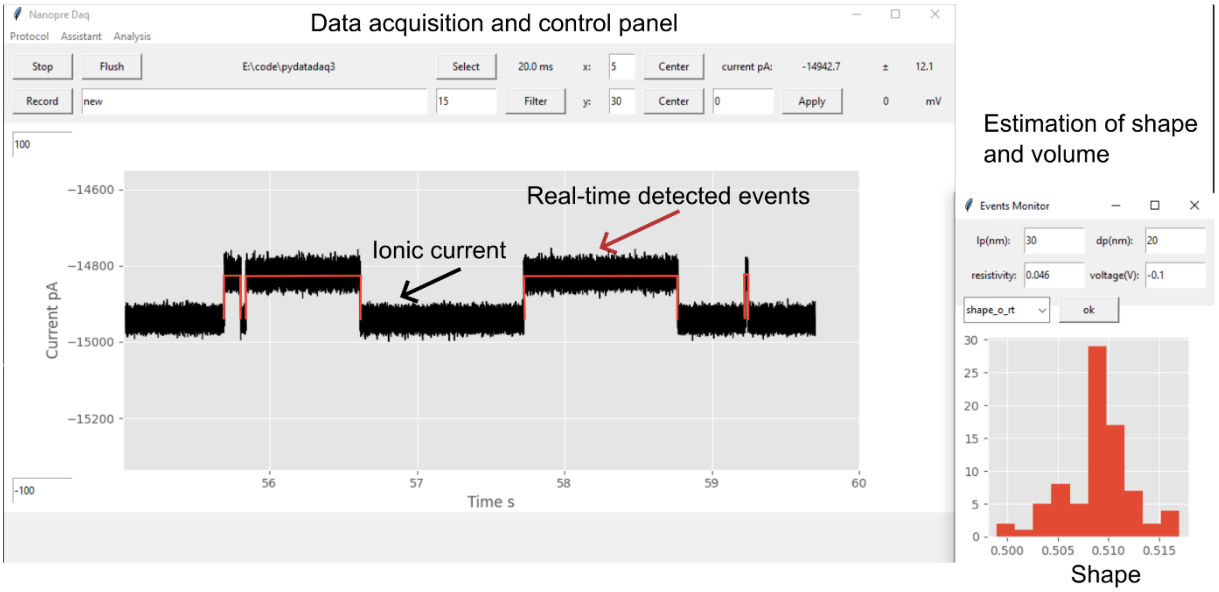


**Figure S1. Screenshot of the data acquisition and real-time estimation of protein using PyDAQ.** The graphical user interface controls the data acquisition, low-pass filter, visualization, resistive pulse detection, and protein estimation. The main plot shows the ionic current (in pA) over time, with detected resistive pulses shown as thin red lines in real time. The histogram on the right provides real-time updates that instantly estimate the shape and volume of the detected proteins.


This manuscript has been submitted to Analytical Chemistry.

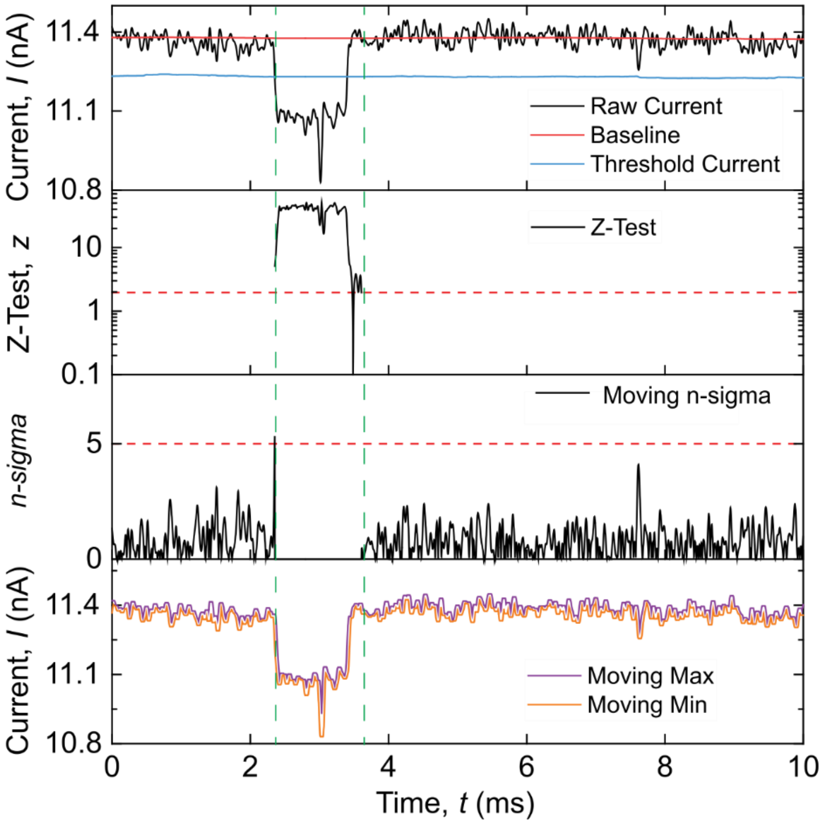


**Figure S2. Example current trace demonstrating the changing of internal variables calculated by the TSW algorithm during the detection of a resistive pulse.** The first panel shows a representative ion current trace containing a single resistive pulse. The blue line represents the continuously updated threshold current for determining the start of the resistive pulse during the TSW processing. The red line represents the moving baseline during the TSW processing. The second panel shows the z-score (*z-test*) to determine the end of the resistive pulse, with the red dashed line indicating a significance threshold of 1.96. The third panel displays the moving *n*-sigma used to determine the start of the resistive pulse, with a threshold of 5 shown by the red dashed line. The bottom panel shows the moving minimum and maximum currents within the statistical window $w_2$.

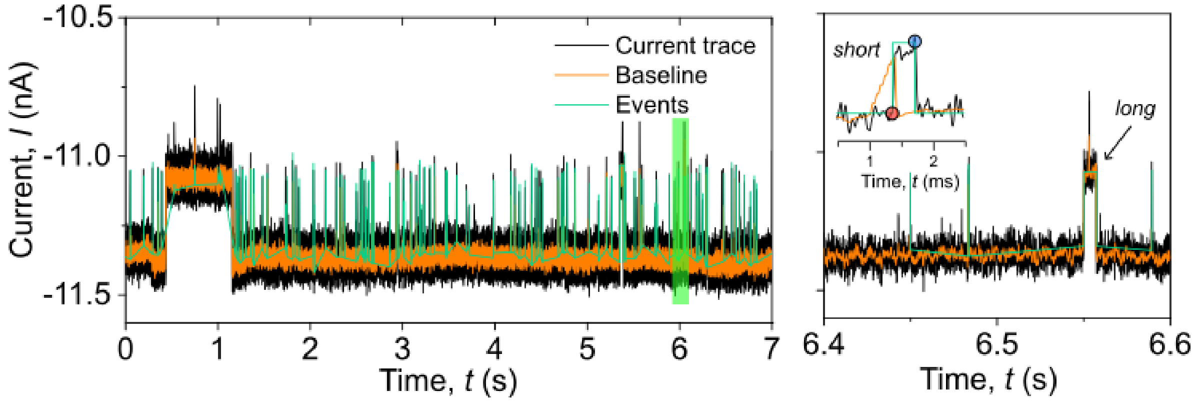


**Figure S3. Performance of the TS peak detection algorithm.** The left panel represents an experimental recording current trace (black line), its baseline (orange line), and resistive pulses (green line) detected using the TS algorithm. The right panel zooms in on the current trace between 5.9 ~ and 6.1 s from the left panel (light green shading). The inset represents a short resistive pulse, its baseline current (orange line), and the start (red circle) and end (blue circle) of the resistive pulses.

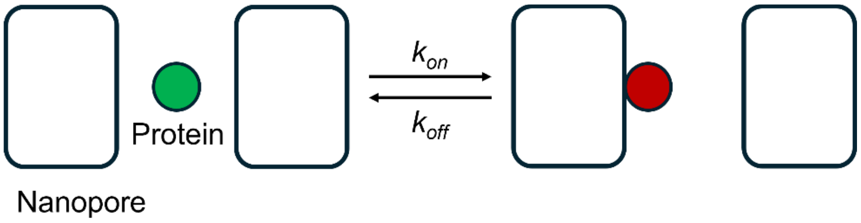


**Figure S4. Schematic illustration of the adsorption of proteins to the wall inside a nanopore.** The green circle represents the proteins that translocate through a nanopore without adsorption, and the red circle represents the proteins that undergo adsorption. The adsorption process can be described as a Markovian reaction.[8] The parameters $k_{on}$ and $k_{off}$ represent stochastic rate constants of the adsorption or dissociation between protein and nanopore surface.

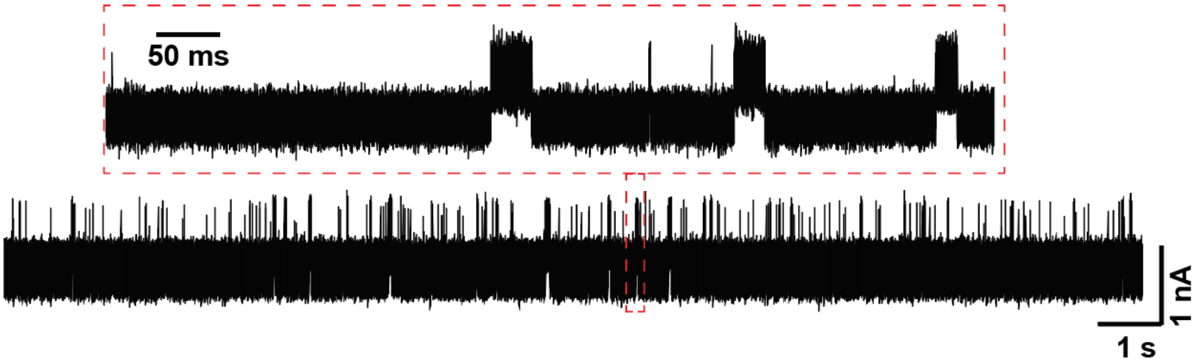



This manuscript has been submitted to Analytical Chemistry.

**Figure S5. Example of simulated current traces.** The trace represents 20 seconds of simulated translocations of oblate particles through a nanopore with a diameter of 20 nm and a length of 30 nm. The red dashed box represents a zoom-in current trace, showing either long resistive pulses from simulated adsorption or short resistive pulses from free translocation of proteins.

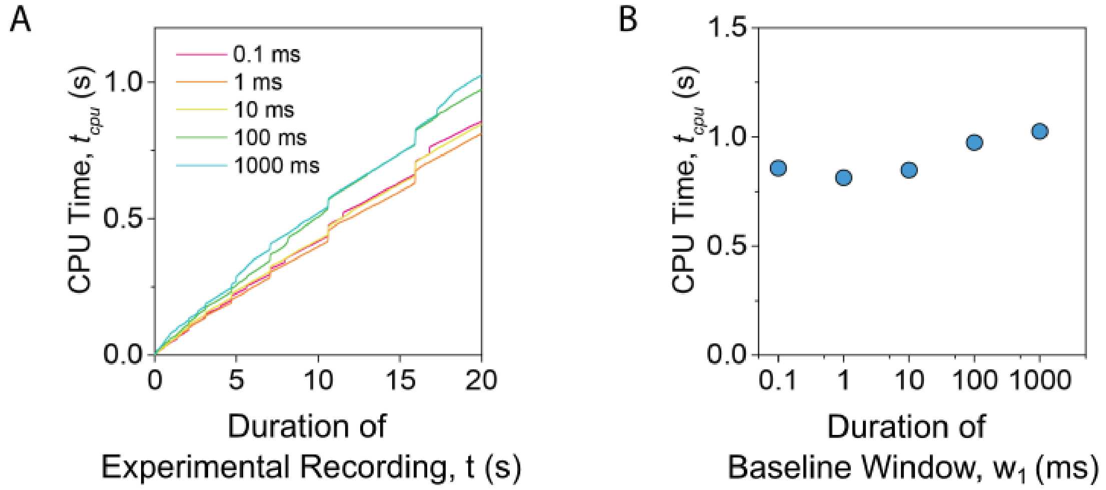


**Figure S6. Analysis of the required time for the TSW algorithm. A.** Cumulative CPU computation time as a function of duration of experimental recording. Colored curves represent the analysis using different durations of the baseline window, $w_1$. **B.** Cumulative CPU time as a function of the duration of the baseline window $w_1$ for analyzing 20 s data at 500 kHz sampling rate. The CPU time does not show significant changes over the size of the baseline window $w_1$.

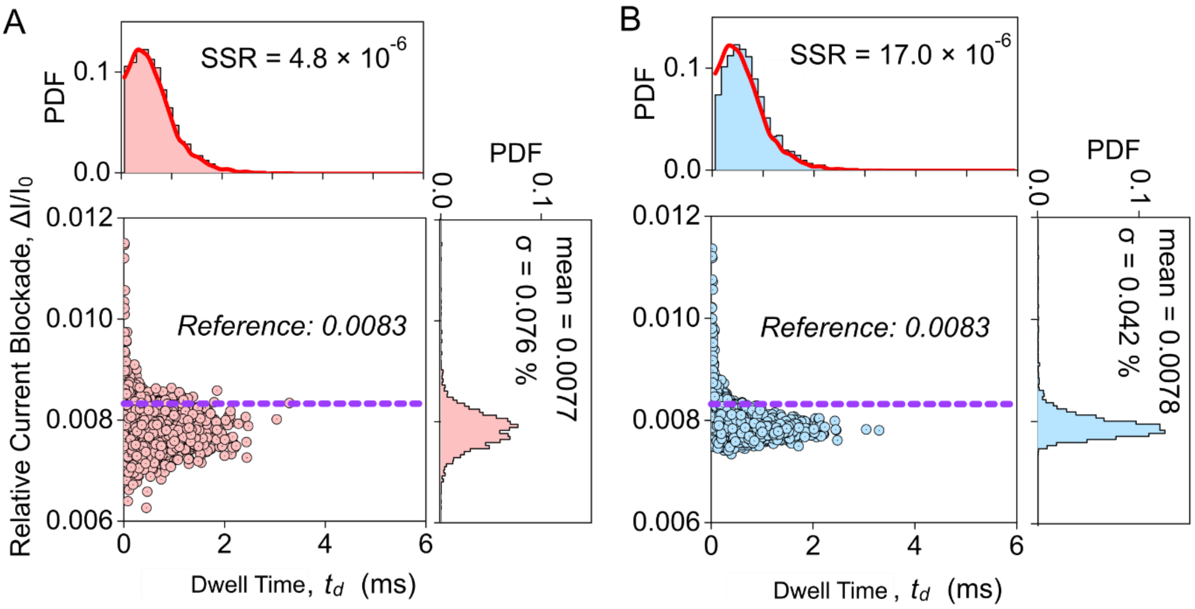


**Figure S7. Comparison of the dwell time and relative current blockades determined by the TS algorithm (A) and the TSW algorithm (B) using simulated data (Supplementary Note 2)**. Each panel shows a 2D scatter



This manuscript has been submitted to Analytical Chemistry.

plot of pulse relative current blockade and dwell time, with marginal histograms for each parameter. The purple dashed line in each scatter plot marks the reference value of the relative current blockades (0.0083). The red curves overlaid on the histograms of dwell time represent the ground truth probability density function (PDF) used in the simulation. The sum of squared residuals (SSR) shows the agreement between the ground truth and the determined dwell time by the TS or TSW algorithm.
